# *YGR042W/MTE1* Functions in Double-Strand Break Repair with *MPH1*

**DOI:** 10.1101/032581

**Authors:** Askar Yimit, TaeHung Kim, Ranjith P. Anand, Sarah Meister, Jiongwen Ou, James E. Haber, Zhaolei Zhang, Grant W. Brown

## Abstract

Double-strand DNA breaks occur upon exposure of cells to agents such as ionizing radiation and ultraviolet light or indirectly through replication fork collapse at DNA damage sites. If left unrepaired double-strand breaks can cause genome instability and cell death. In response to DNA damage, proteins involved in double-strand break repair by homologous recombination re-localize into discrete nuclear foci. We identified 29 proteins that co-localize with the recombination repair protein Rad52 in response to DNA damage. Of particular interest, Ygr042w/Mte1, a protein of unknown function, showed robust colocalization with Rad52. Mte1 foci fail to form when the DNA helicase Mph1 is absent. Mte1 and Mph1 form a complex, and are recruited to double-strand breaks *in vivo* in a mutually dependent manner. Mte1 is important for resolution of Rad52 foci during double-strand break repair, and for suppressing break-induced replication. Together our data indicate that Mte1 functions with Mph1 in double-strand break repair.

---

Effective repair of double-strand DNA breaks (DSBs) is critical to the preservation of genome stability, yet most modes of DSB repair have significant potential to generate sequence alterations or sequence loss. Repair of DSBs by homologous recombination can result in loss of heterozygosity when resolution of recombination intermediates between homologous chromosomes results in a crossover. As such, cells possess several mechanisms by which crossing over can be suppressed in favor of non-crossover recombination products. Double Holliday junction intermediates that result from invasion of a homologous chromosome by both ends of a resected DSB (Szostak *et al*. 1983) can be resolved nucleolytically, by the action of the Yen1 and Mus81/Mms4 endonucleases (Blanco *et al*. 2010; Ho *et al*. 2010), to produce a random distribution of crossover and noncrossover products. By contrast, the same dHJ intermediates can be dissolved by the combined helicase and ssDNA decatenase action of the BTR complex (Sgs1/Top3/Rmi1 in yeast) (Wu *et al*. 2006; Yang *et al*. 2010) to yield exclusively non-crossover products (Wu and Hickson 2003). Crossovers can also be prevented if the D-loop structure that results from the first strand invasion by one end of a resected DSB into the homologous chromosome is unwound before capture of the second end to form the dHJ. Unwinding of D-loops is catalyzed *in vitro* and *in vivo* by the 3ʹ to 5ʹ DNA helicase Mph1 (Prakash *et al*. 2009; Sun *et al*. 2008) in order to prevent loss of heterozygosity due to crossovers and break-induced replication (Luke-Glaser and Luke 2012; Mazon and Symington 2013; Stafa *et al*. 2014).

The Mph1 DNA helicase was first identified as a deletion mutant with an increased mutation frequency (Entian *et al*. 1999). Subsequent characterization revealed that *mph1* mutants are sensitive to the alkylating agent MMS, and to a lesser degree to ionizing radiation (Scheller *et al*. 2000), and that *mph1* mutants are proficient for mitotic recombination (Schurer *et al*. 2004). Molecular insight into Mph1 function in recombination reactions comes from evidence that Mph1 is a DNA helicase (Prakash *et al*. 2005), and that Mph1 can unwind Rad51 D-loops (Prakash *et al*. 2009; Sun *et al*. 2008) and extended D-loops (Sebesta *et al*. 2011). Consistent with an anti-recombination role for Mph1, overexpression of *MPH1* reduces recombination rate and reduces loading of Rad51 at an induced DSB (Banerjee *et al*. 2008). Indeed, Mph1 suppresses crossing over during mitotic recombination, likely by unwinding D-loop recombination intermediates formed by Rad51 (Prakash *et al*. 2009) and preventing ectopic resolution of early strand exchange intermediates by the Mus81-Mms4 nuclease (Mazon and Symington 2013). Mph1 inhibits break-induced replication (BIR) repair of double-strand breaks (Luke-Glaser and Luke 2012) and promotes template switching during BIR (Stafa *et al*. 2014), both consistent with the ability of Mph1 to unwind recombination intermediates *in vitro*. In addition to functioning in crossover suppression, Mph1 plays a pro-recombinogenic role in repair of stressed DNA replication forks (Chavez *et al*. 2011; Chen *et al*. 2009 2013; Choi *et al*. 2010; Sun *et al*. 2008; Xue *et al*. 2014; Zheng *et al*. 2011), and inhibits non-homologous end-joining repair at telomeres (Luke-Glaser and Luke 2012). Mph1 is thought to be the functional homologue of the human FANCM protein. Thus, available evidence points to diverse functions for Mph1, and these functions are likely connected to the ability of Mph1 to unwind and remodel DNA structures.

Here we leverage intracellular protein location data to identify the complement of proteins that co-localize with the recombination repair protein Rad52 in nuclear foci during the response to DNA double-strand breaks. In addition to defining the membership of Rad52 foci, we identify an uncharacterized protein, Ygr042w/Mte1, that functions in double-strand break repair. Mte1 acts in complex with Mph1 at double-strand breaks *in vivo*, is important for DSB repair as assessed by resolution of Rad52 foci, and functions, as is the case for Mph1, in suppressing break-induced replication repair of double-strand DNA breaks.

## Materials and Methods

### Yeast strains and media

All yeast strains used in this study are derivatives of BY4741 (Brachmann *et al*. 1998), CL11–7, or W303, and are listed in Table S1. Strains were constructed using genetic crosses and standard PCR-based gene disruption techniques. Standard yeast media and growth conditions were used.

### Chromatin immunoprecipitation and deep sequencing

Chromatin immunoprecipitation was performed using Flag-epitope tagged versions of each indicated protein, as previously described (Balint *et al*. 2015; Roberts *et al*. 2008), with modifications. Cells were grown to mid-logarithmic phase in YPR (1% yeast extract, 2% peptone, 3% raffinose) at 28°C and then arrested in G2/M with 20 *μ*g/ml nocodazole for 4 hrs. Galactose was added to 2% final to induce expression of the HO endonuclease. Cells were sampled before galactose addition and after 4 hours of induction, and cross-linked with formaldehyde overnight. Cells were harvested and washed twice with cold TBS (20 mM Tris-HCl pH 7.5, 150 mM NaCl), resuspended in FA-lysis buffer (50 mM HEPES pH 7.5, 2 mM EDTA, 1% Triton X-100, 0.1% sodium deoxycholate, 150 mM NaCl) containing 0.05% SDS, lysed and sonicated. Immunoprecipitates were washed sequentially with 1 ml of FA-lysis buffer, FA-lysis buffer containing 1M NaCl, FA-lysis buffer containing 0.5M NaCl, Wash Buffer (50 mM HEPES pH 7.5, 0.25M LiCl, 2 mM EDTA, 1% Triton X-100, 1% sodium deoxycholate, 1% NP-40,10 mM Tris-HCl pH 8.0), and TE (10 mM Tris-HCl pH 8.0, 1 mM EDTA). Protein-DNA complexes were eluted, cross-links were reversed, protein and RNA was digested, and DNA was isolated by phenol/chloroform extraction and ethanol precipitation. Sequencing libraries were generated using the Nextera XT DNA Sample Preparation Kit (Illumina) with custom index primers for the PCR amplification step. Libraries were quantified using a 2100 Bioanalyzer (Agilent) and the KAPA SYBR FAST Universal qPCR Kit (KAPA Biosystems).

### Sequencing data analysis

Input and IP samples from each experiment were sequenced on an Illumina HiSeq 2500 (50 nucleotide singleend reads). All sequencing data are deposited in the Sequence Read Archive (http://www.ncbi.nlm.nih.gov/sra, study accession SRP064493). The number of reads for each sample ranges from 12.8 M – 25.7 M. The quality of sequencing reads was first assessed using FastQC. (http://www.bioinformatics.bbsrc.ac.uk/projects/fastqc). All samples have a median PHRED score of 30 or higher for all positions. Sequenced reads were mapped to the *S. cerevisiae* reference genome version WS220 (downloaded from the Saccharomyces Genome Database (Cherry *et al*. 2012; Engel *et al*. 2014)) using Bowtie2 (version 2.0.0) (Langmead and Salzberg 2012) with default settings, except for forcing end-to-end alignment. Greater than 96% mapping rates were achieved for all samples, yielding a minimum 50x coverage for all samples (Table S2). In order to reduce any bias from DNA sequencing, the data were normalized by the ratio of coverage for each IP and input pair prior to each comparison. We used a 100 bp sliding window with a step size of 50 bp to calculate enrichment scores as a log2 ratio of normalized read counts for each IP:input pair. For the enrichment scores displayed in Figure 5, the enrichment score for each of the 0h samples was subtracted from each of the matched 4h samples. Figure S1 displays the enrichment scores for all of the IP:input pairs.

### Whole cell extracts, immunoblotting and immunoprecipitation

Logarithmically growing cells at 30°C were treated with or without 5 *μ*g/ml phleomycin (BioShop PEO422.25) for 2 hours before cells were collected, fixed with 10% trichloroacetic acid, and whole cell extracts were prepared (Pellicioli *et al*. 1999). Proteins were resolved by SDS-PAGE and subjected to immunoblotting with *α*-Flag M2 (F3165, Sigma-Aldrich), *α*-HA (ab16918, Abcam), or *α*-tubulin (YOL1/34, Serotec) antibodies. Native extracts for immunoprecipitation were prepared from 5×10^8^ cells as previously described (Shimomura *et al*. 1998), with some modifications. Cell pellets were resuspended in FA-lysis buffer containing 1 mM DTT, 2 mM sodium fluoride, 1 mM sodium ortho-vanadate, 1X Complete Mini EDTA-free protease inhibitor cocktail (Roche 11836170001), 2.5 *μ*g/ml aprotinin, 10 mM *β-*glycerophosphate, 5 *μ*g/mL leupeptin, 2 *μ*g/mL pepstatin A, 1 mM PMSF, and 5 *μ*g/ml TLCK, then lysed with glass beads. Cleared extracts were immunoprecipitated with *α*-Flag M2 antibody. Beads were washed twice with 0.5 ml FA-lysis buffer as above, and eluted in 5X SDS loading buffer.

### DNA damage sensitivity

Yeast strains were grown overnight in YPD, diluted serially, and spotted onto YPD plates containing the indicated concentrations of phleomycin. Plates were incubated at 30°C for 2–3 days before imaging.

### Fluorescence microscopy

For analysis of GFP fusion protein nuclear foci, strains were grown to mid-log phase in YPD, diluted into fresh YPD and cultured overnight to OD_600_ = 0.3. Cells were treated for 120 minutes with 5 *μ*g/ml phleomycin, or cultured without phleomycin, harvested, and washed once in low fluorescence medium with or without phleomycin before imaging. Eleven z slices with a 0.4 *μm* step size were acquired using Volocity imaging software (PerkinElmer) controlling a Leica DMI6000 confocal fluorescence microscope with fluorescein isothiocyanate, Texas Red and differential interference contrast filter sets (Quorum Technologies). Images were scored by visual inspection for GFP fusion protein foci. Samples were compared using the t-test or the Wilcoxon rank sum test, as appropriate, in R (www.r-project.org). Data were plotted using ggplot2, in R. For Rad52-GFP foci, the same procedure was used except that cells were blocked in G2/M phase by treatment with 20 *μ*g/ml nocodazole for 3 hours, and exposed to 50 *μ*g/ml phleomycin for 30 minutes.

### Recombination assays

Recombination rates were calculated using a direct repeat recombination assay (Smith and Rothstein 1999) and quantifying recombination from the number of Leu+ recombinant colonies using the method of the median (Lea and Coulson 1949). Each fluctuation test comprised 9 independent cultures, and the results from 10 fluctuation tests were plotted in R. Rates were compared using a Welch two-sample t-test in R.

BIR efficiencies were calculated as described previously (Anand *et al*. 2014). Briefly, cells were plated for individual colonies on YEPD + clonNat to retain the HOcs (which is marked with *natMX*). Approximately one million cells from individual colonies were appropriately diluted and plated on YEPD plates to get the total cell count and on YEP-Gal plates for HO induction. Cells that grew on YEP-Gal plates (DNA break-survivors) were counted and replica plated to plates lacking uracil to determine BIR frequencies. For each replicate, Ura+ frequencies were calculated as the total Ura+ cells that grew on plates lacking uracil over the total cells on YEPD. Experiments were repeated at least 3 times, plotted in R, and compared using a Welch two-sample t-test in R.

### Data Availability

Strains are available upon request. Table S1 contains the genotypes of all strains used. Table S2 contains statistics for all deep sequencing, including NCBI Sequence Read Archive (SRA) accession numbers.

## Results

### Twenty-nine proteins form nuclear foci that detectably co-localize with Rad52 foci

A number of DNA repair proteins change their intracellular localization from pan-nuclear to nuclear foci in response to DNA damage. Proteins that localize in nuclear foci have been identified in candidate approaches (Burgess *et al*. 2009; Germann *et al*. 2011; Lisby *et al*. 2004 2001; Melo *et al*. 2001; Zhu *et al*. 2008) and in genome-scale screens (Denervaud *et al*. 2013; Mazumder *et al*. 2013; Tkach *et al*. 2012; Yu *et al*. 2013). Nuclear foci are commonly thought of as centers of DNA repair, in part because foci formed by recombination repair proteins co-localize with double-strand DNA breaks (Lisby *et al*. 2003). However, not all nuclear foci are identical to the canonical DNA repair centers that are marked by the recombination protein Rad52. For example, Cmr1 forms foci that do not co-localize detectably with Rad52 (Tkach *et al*. 2012), but rather co-localize with a distinct set of proteins in an intranuclear quality control compartment (Gallina *et al*. 2015).

We tested 61 budding yeast proteins that form nuclear foci in response to DNA damage to identify those that co-localize detectably with Rad52. Nuclear foci proteins were tagged with GFP (Huh *et al*. 2003), Rad52 was tagged with mCherry, and cells were examined by fluorescence microscopy after treatment with the double-strand DNA break inducing agent phleomycin (Figure 1). Twenty-nine proteins co-localized detectably with Rad52 (Figure 1A and Tables S3, S4, and S5). The extent of co-localization ranged from 79% of foci for Srs2, to 2% of foci for Csm1 (Table S3). Fourteen proteins had not previously been described as components of Rad52 foci (Figure 1A and 1C), although most are known DNA repair, DNA replication, or checkpoint signaling proteins (Figure 1D). We identified one protein, Ygr042w, with no known role in recombination repair. Mutants in *YGR042W* affect telomere length (Askree *et al*. 2004), and the fission yeast homologue of Ygr042w, Dbl2, forms foci that co-localize with an induced double-strand DNA break (Yu *et al*. 2013). The extensive co-localization of Ygr042w with Rad52 foci, similar to the extent of co-localization observed for members of the Rad52 epistasis group (Symington 2002) Rad55, Rad57, and Rad59, suggests that Ygr042w could function in repair of double-strand DNA breaks. While this work was in progress, a name for *YGR042W* was reserved in the Saccharomyces Genome Database, *MTE1* (Mph1-associated Telomere maintenance protein). Thus, we now refer to *YGR042W* as *MTE1*.

**Figure 1.**
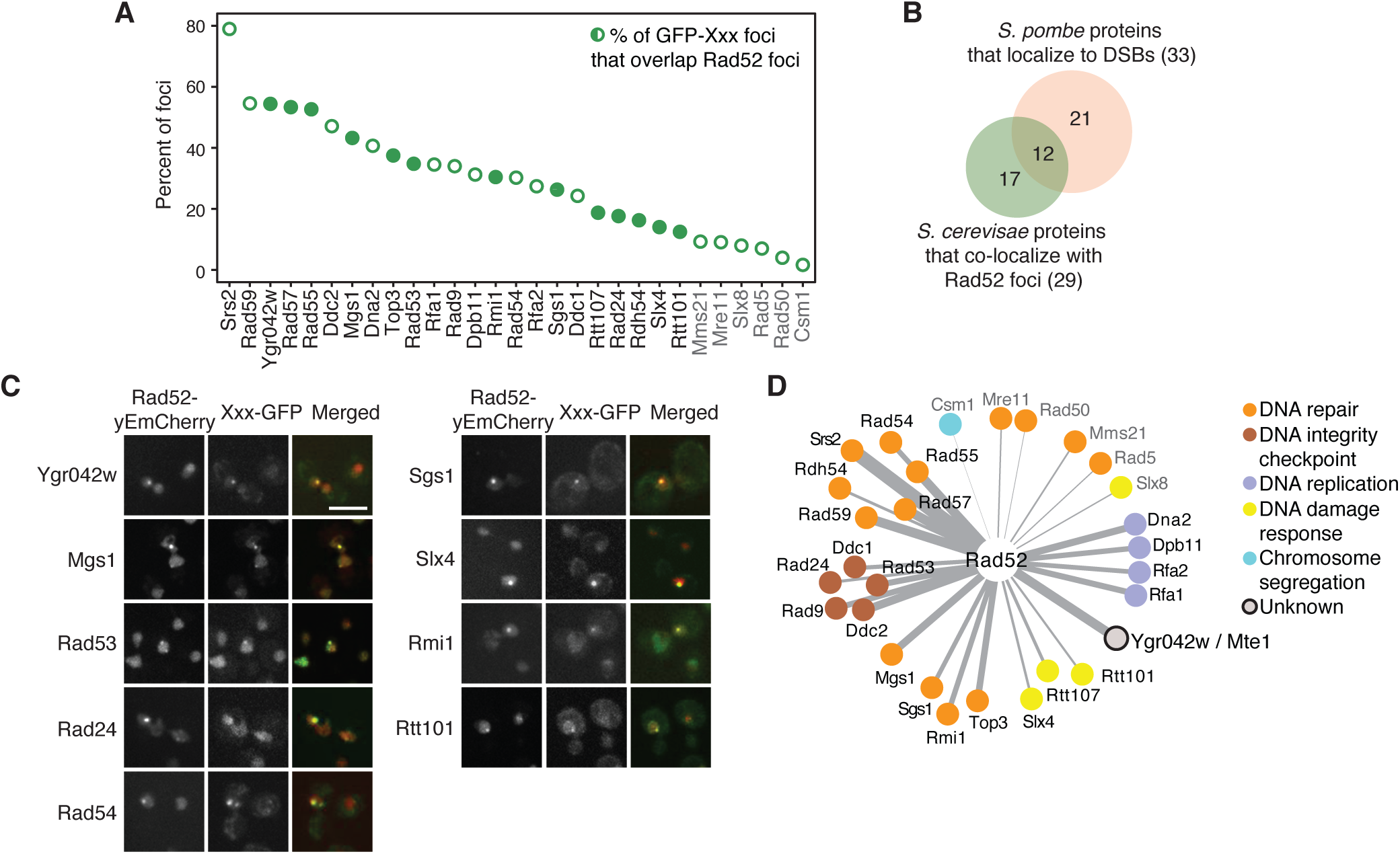
Twenty-nine proteins form nuclear foci that detectably co-localize with Rad52 foci. (A) The percent of nuclear foci formed by each GFP fusion protein that overlap with Rad52-yEmCherry foci after 2 hours in 5 *μ*g/ml phleomycin is plotted. Open circles indicate co-localizations that were previously identified. Closed circles indicate Rad52 co-localizations that have not been previously described. Protein names in grey indicate those with a percent co-localization at or below that seen with Mre11. (B) The overlap between the proteins that co-localize with Rad52 foci and those that co-localize with an induced double-strand DNA break in fission yeast is shown. (C) Representative fluorescence micrographs showing co-localization of the indicated GFP fusion proteins with Rad52 foci. The mCherry, GFP, and merged images are shown. The scale bar is 5 *μm*. (D) A network of the proteins that co-localize detectably with Rad52. Protein function is indicated by colour and edge thickness is proportional to the extent of protein co-localization with Rad52 foci.

### Mte1 foci form in S/G2 phase and in response to double-strand breaks

The foci formed by Mte1 in response to phleomycin localize to the nucleus (Figure 2A) and form more frequently in cells in S and G2 phases than in G1 cells (Figure 2B). Mte1 foci also form in the absence of DNA damaging agents, in 13% of cells during S or G2 phase, but in only 3% of cells during G1 phase (Figure 2B), similar to Rad52 foci (Lisby *et al*. 2001). As expected, Mte1 foci levels increase with increasing phleomycin concentration and with increasing time of phleomycin exposure (Figure 2C). Deletion of *MTE1* confers modest sensitivity to phleomycin, but not to other DNA damaging and replication stress agents, methyl methanesulfonate, hydroxyurea, and camptothecin (Figure 2D).

**Figure 2.**
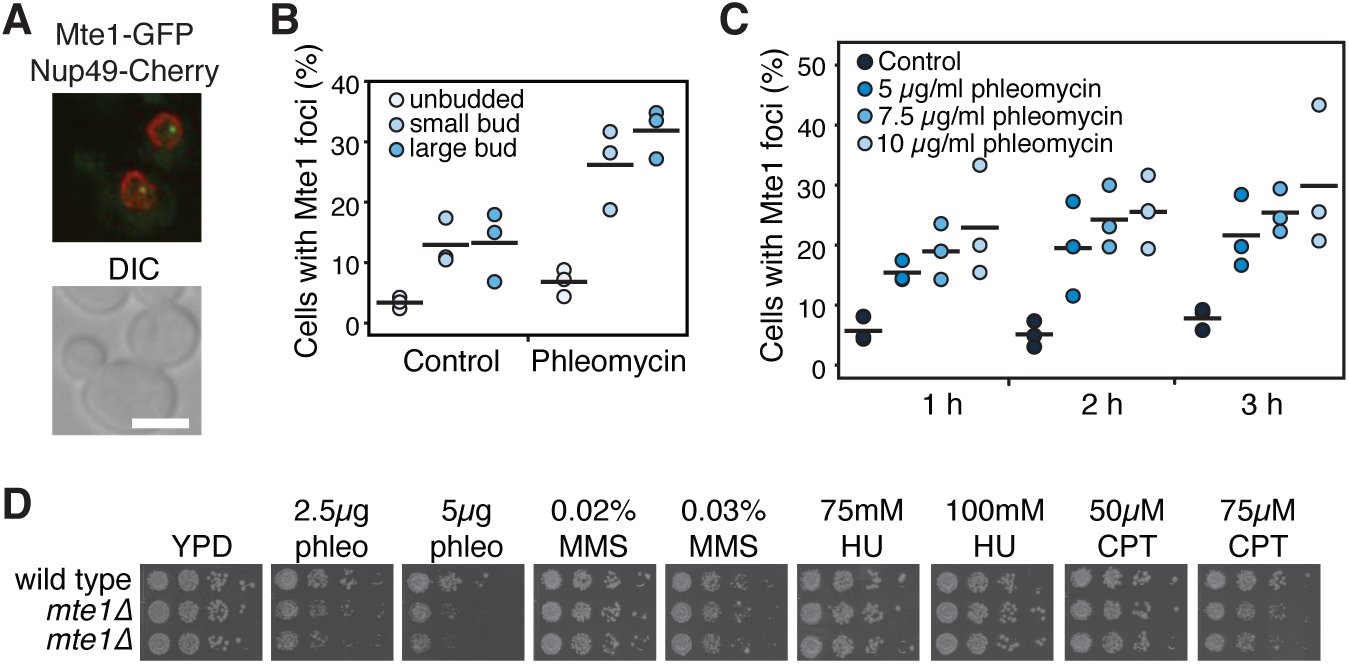
Mte1 foci form in S/G2 phase and in response to double strand breaks. (A) Mte1-GFP nuclear foci are shown in a merged fluorescence micrograph. The lower panel is the DIC image of the same cells. The scale bar is 5 *μ*m. (B) The fraction of unbudded, small-budded, and large-budded cells with Mte1 nuclear foci after 2 hours in 5 *μ*g/ml phleomycin is plotted. The black bars show the means of the 3 replicates. (C) The fraction of cells with Mte1 nuclear foci after 1, 2, or 3 hours in the indicated concentrations of phleomycin is plotted. The black bars show the means of the 3 replicates. (D) Serial ten-fold dilutions of the indicated strains were spotted on the indicated concentrations of phleomycin (phleo), methyl methanesulfonate (MMS), hudroxyurea (HU), or camptothecan (CPT). Plates were photographed after 2 to 3 days.

### Mte1 foci are increased when end-resection is defective and depend on MPH1

We tested whether Mte1 focus formation was altered in mutants of genes encoding other nuclear focus proteins and several additional DNA repair proteins. Of the 52 mutants tested, 5 led to increased Mte1 focus formation (Figure 3A and Table S6). Three of the mutants, in *MRE11, RAD50*, and *XRS2*, would eliminate the DSB end-resection function of the MRX complex (Ivanov *et al*. 1994), and *RAD52* is critical for formation of the Rad51 filament at resected DSBs (Sugawara *et al*. 2003), among other functions. Mte1 foci increase in both the presence and absence of phleomycin in *mre11*Δ, *rad50*Δ, *xrs2*Δ, and *rad52*Δ, indicating that spontaneous DSBs are either more prevalent in these mutants, or are repaired less effectively. By contrast, the *rad9*Δ mutant, which is defective in DNA damage checkpoint signaling and results in faster end-resection at a double-strand break (Ferrari *et al*. 2015; Lazzaro *et al*. 2008), displays increased Mte1 foci only in the presence of phleomycin. We tested whether other checkpoint mutants result in increased Mte1 foci (Figure 3A). We disrupted checkpoint signaling upstream of Rad9, by deleting *MEC1, TEL1*, or both, and found that only the *mec1*Δ *tel1*Δ double mutant had a statistically evident increase in Mte1 foci, in both the absence and presence of phleomycin (p=5.2×10^−5^ and p=0.00095, one-sided t-test). Interestingly, *mec1*Δ *tel1*Δ cells, like *rad9*Δ, have a higher rate of resection (Tsabar *et al*. 2015), and so increased Mte1 foci in these mutants could reflect increased resection of the DSB.

**Figure 3.**
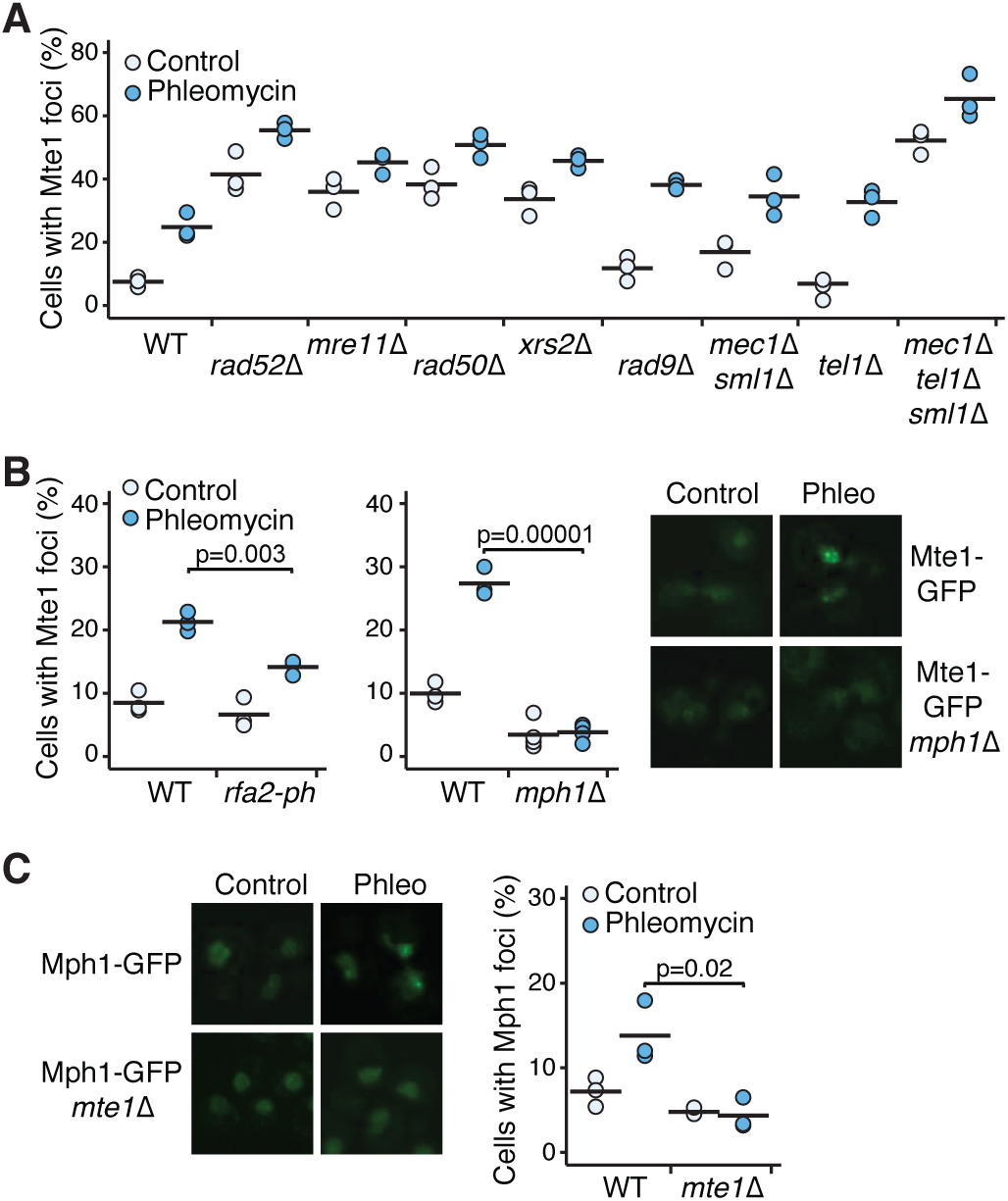
Mte1 foci are increased in *MRX* mutants and depend on *MPH1*. (A) The percent of cells with Mte1 foci is plotted for mutants with increased numbers of foci. Three replicates for each strain, in the absence and presence of 5 *μ*g/ml phleomycin, are plotted. The black bars show the means of the 3 replicates. N ranged from 41 to 211 cells per strain per replicate. (B) The percent of cells with Mte1 foci is plotted for mutants with decreased numbers of foci (left), for untreated cells and cells grown in the presence of 5 *μ*g/ml phleomycin for 2 hours. The black bars show the means of the replicates. The indicated samples were compared using a one-sided t-test. N ranged from 65 to 174 cells per strain per replicate. Representative images (right) of cells with Mte1 foci are shown for untreated cells and cells grown in the presence of 5 *μ*g/ml phleomycin for 2 hours, for wild type cells and *mph1*Δ cells. (C) Representative images (left) and the percent of cells with Mph1 foci (right) are shown for untreated cells, and cells grown in the presence of 5 *μ*g/ml phleomycin for 2 hours, for wild type cells and *mte1*Δ cells. The black bars show the means of the replicates. The indicated samples were compared using a one-sided t-test. N ranged from 89 to 276 cells per strain per replicate.

Two mutants, *mph1*Δ and *rpa2-ph*, caused decreased Mte1 focus formation (Figure 3B). Mph1 and RPA are proposed to function together to suppress recombination (Banerjee *et al*. 2008). Mph1 forms nuclear foci in unperturbed cells and in MMS (Chen *et al*. 2009), and we find that Mph1 foci increase in the presence of phleomycin (Figure 3C). Deletion of *MTE1* reduces Mph1 foci to background levels (Figure 3C), suggesting that Mph1 and Mte1 might function in concert.

### Mte1 and Mph1 interact physically and are in the same genetic pathway

We tested whether Mte1 interacts with Mph1 in co-immunoprecipitation experiments (Figure 4). We found that Mte1 immunoprecipitates contain Mph1 (Figure 4A) and that Mph1 immunoprecipitates contain Mte1 (Figure 4B). Mte1 and Mph1 appear to interact constitutively, as the extent of co-immunoprecipitation is unaffected by the presence of phleomycin. Consistent with Mte1 and Mph1 forming a complex, 38% of Mte1 foci co-localize with Mph1 after 3h in phleomycin (Figure 4C). Both *mte1*Δ and *mph1*Δ confer modest sensitivity to phleomycin, and the double mutant *mte1*Δ *mph1*Δ is no more sensitive than either of the single mutants, suggesting the *MTE1* and *MPH1* function in the same genetic DSB response pathway. By contrast, *mte1*Δ and *rad52*Δ show additive phleomycin sensitivity (Figure 4D and 4E) suggesting that *MTE1* and *RAD52* play non-redundant roles in DSB repair.

**Figure 4.**
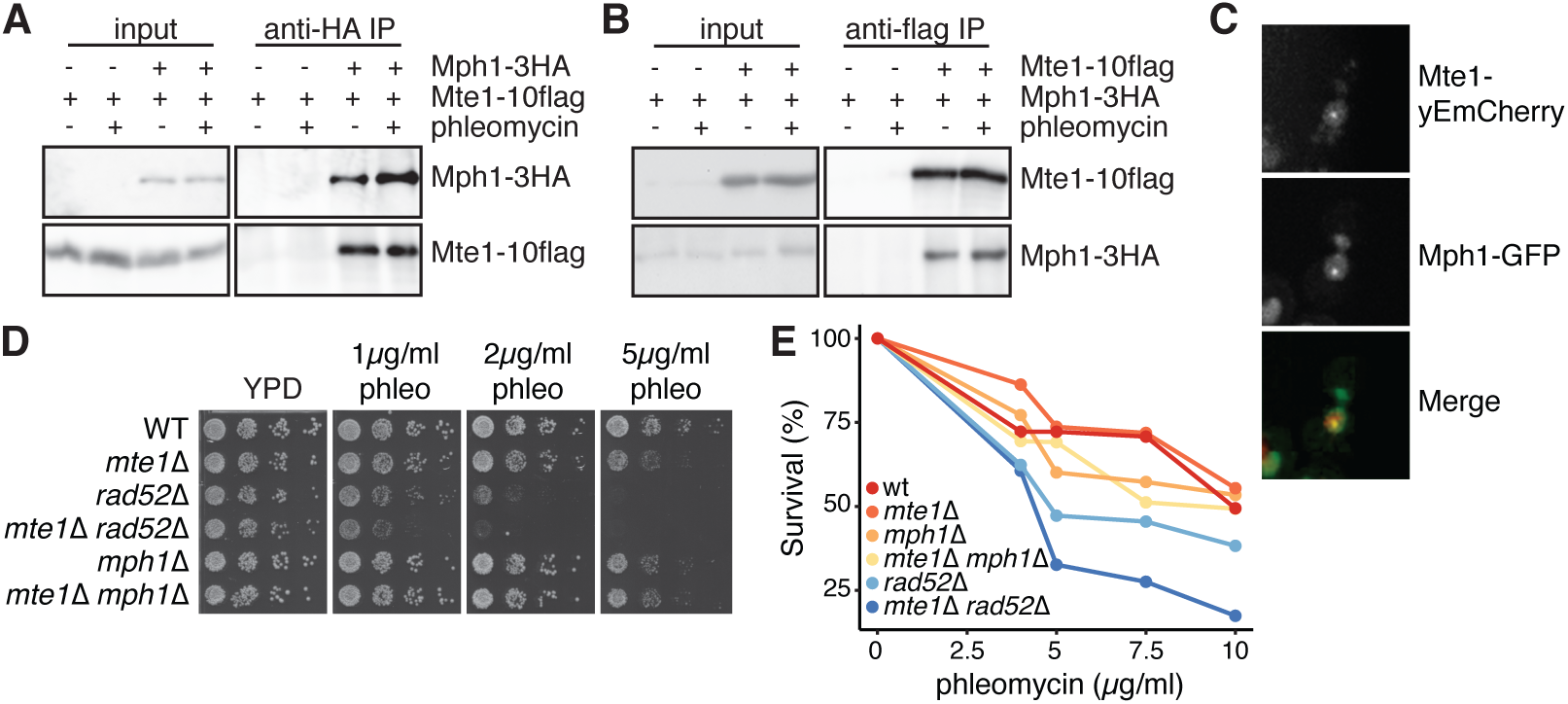
Mte1 and Mph1 interact physically and are in the same genetic pathway. (A) Extracts of cells expressing Mte1–10flag and Mph1–3HA proteins as indicated were subjected to immunoprecipitation with an anti-HA antibody. Input and immunoprecipitate (IP) fractions were immunoblotted to detect Mte1–10flag or Mph1–3HA. (B) In the reciprocal of panel A, extracts of cells expressing Mte1–10flag and Mph1–3HA proteins as indicated were subjected to immunoprecipitation with an anti-flag antibody. Input and immunoprecipitate (IP) fractions were immunoblotted to detect Mte1–10flag or Mph1–3HA. (C) Representative fluorescence micrographs showing co-localization of Mte1 with Mph1 following phleomycin treatment. The mCherry, GFP, and merged images are shown. (D) Serial ten-fold dilutions of the indicated strains were spotted on media containing the indicated concentrations of phleomycin. Plates were photographed after 2 to 3 days. (E) The indicated strains were cultured in the presence of the indicated concentrations of phleomycin for 2 hours, diluted and plated on media lacking phleomycin. The fraction of cells that formed colonies is plotted.

### Mte1 and Mph1 localize to double-strand DNA breaks

Many proteins involved in double-strand DNA break repair are physically associated with chromatin adjacent to strand breaks *in vivo*, including Mph1 (Prakash *et al*. 2009). We used chromatin immunoprecipitation followed by deep sequencing to assess the binding of Mte1 and Mph1 to the region flanking an induced HO double-strand break (Figure 5). The HO double-strand break was induced by growth in galactose to induce expression of the HO endonuclease. Cultures were sampled before HO induction, and after 4 hours in galactose, cross-linked with formaldehyde, and subjected to chromatin immunoprecipitation. The enrichment of DNA sequences in the immunoprecipitate relative to the input sample indicates regions of protein binding. We first tested Rad52, which is known to localize robustly to DSBs *in vivo* (Wolner *et al*. 2003), and found a peak of enrichment on chromosome III following HO induction, centered on the HO endonuclease site, (Figure 5A and Figure S1). Similar peaks were detected at the induced DSB for both Mte1 and Mph1, indicating that the Mte1-Mph1 protein complex is recruited to DNA double-strand breaks *in vivo* (Figure 5A and S1). Of particular interest, the Mte1 enrichment at the DSB was reduced in an Mph1Δ mutant, and Mph1 enrichment at the DSB was reduced in an *mte1*Δ mutant (Figure 5A and S1). Mte1 and Mph1 protein levels were unchanged in the mutant backgrounds (Figure 5B), suggesting that the functional unit recruited to DSBs is an Mte1-Mph1 complex.

**Figure 5.**
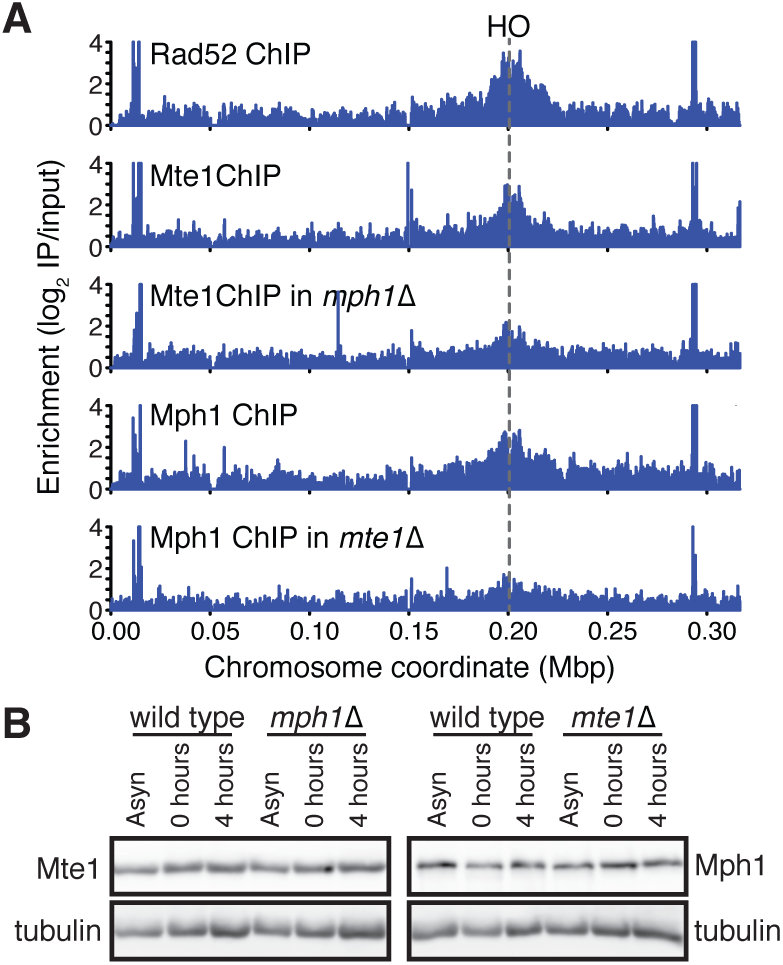
Mte1 and Mph1 are recruited to double-strand DNA breaks. (A) ChIP-seq analysis was performed on *RAD52-FLAG, MTE1-FLAG, MTE1-FLAG mph1*Δ*, MPH1-FLAG*, and *MPH1-FLAG mte1*Δ cells at 0h and 4h following the induction of a specific double-strand break at the *MAT* locus by the HO endonuclease. ChIP enrichment scores derived by subtracting the log2 immunoprecipitate to input ratio at 0h from the ratio at 4h are plotted across chromosome III. The position of the HO cut site is indicated. (B) Extracts from cells used in panel A were subjected to immunoblot analysis, and probed with an anti-flag antibody and an anti-tubulin antibody (as a loading control).

### Increased phleomycin-induced DSBs in the absence of Mte1

The presence of Mte1 at an induced DSB, and the sensitivity of *mte1*Δ strains to DSBs, suggested that Mte1 could play a role in DSB repair. We measured Rad52 focus formation as a proxy for the presence of DNA damage. Cells were blocked in G2 phase with nocodazole and treated with 50 *μ*g/ml phleomycin for 30 minutes. Phleomycin caused an increase in the fraction of cells with Rad52 foci in *mph1*Δ, *mte1*Δ, and the *mph1*Δ *mte1*Δ double mutant compared to the wild-type (Figure 6A), and an increase in Rad52 focus intensity (Figure 6B). The *mph1*Δ and *mte1*Δ single mutants and the *mph1*Δ *mte1*Δ double mutant had similar effects in both assays, suggesting that Mte1 and Mph1 function together in DSB repair. We measured recombination directly in *mte1*Δ mutants (Figure 6C). In the absence of DNA damage, *mte1*Δ, like *mph1*Δ (Schurer *et al*. 2004), is proficient in mitotic recombination, displaying a recombination rate that is highly similar to the wild type.

**Figure 6.**
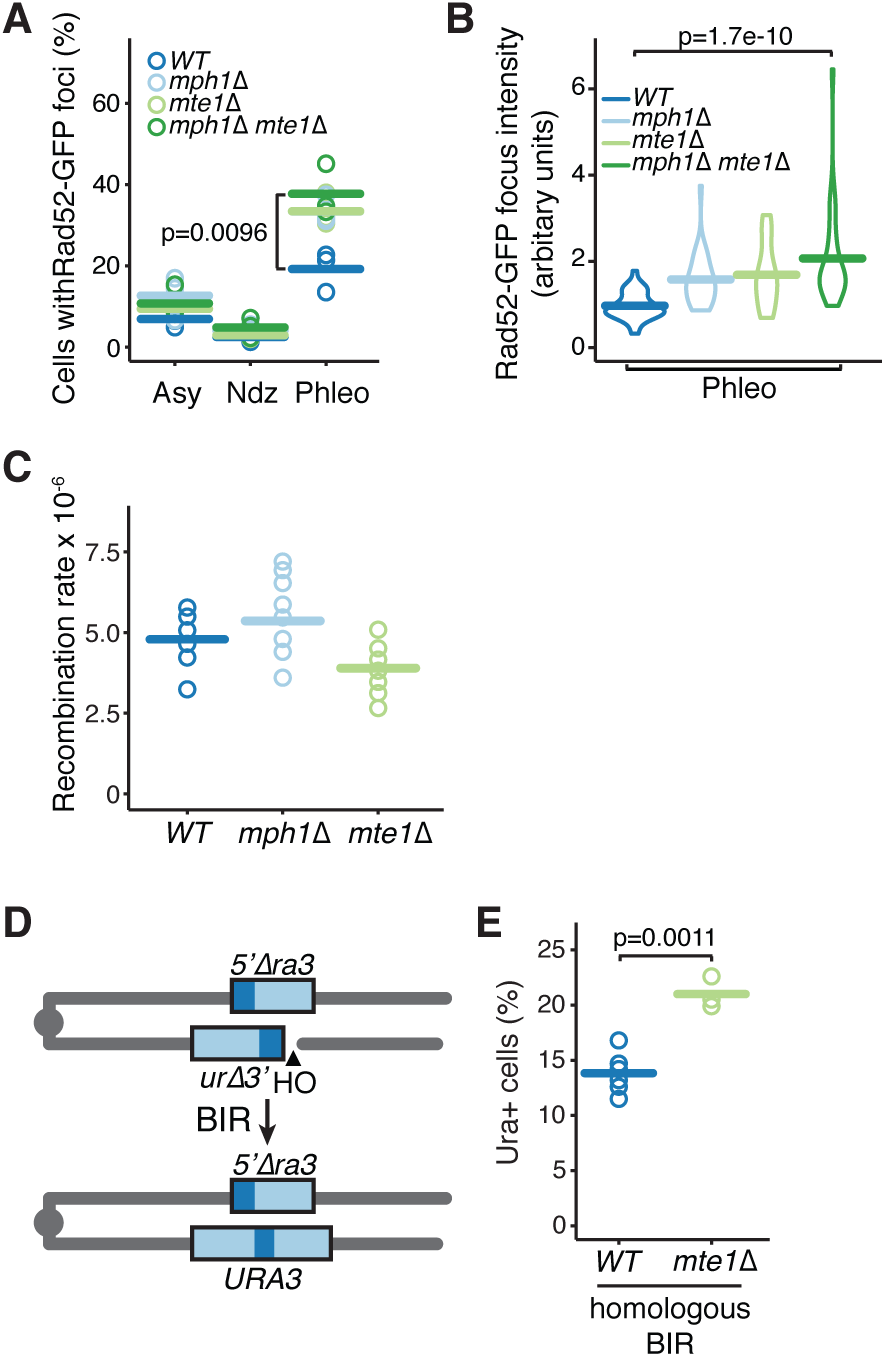
Mte1 contributes to double-strand break repair. (A) The percent of cells with Rad52 foci is plotted for the indicated strains. Samples were from logarithmic phase (Asy), G2/M (Ndz), and after treatment with 50 *μ*g/ml phleomycin for 30 minutes (Phleo). Three replicates for each strain, for each condition, are plotted. The bars show the mean of the 3 replicates. N ranged from 42 to 160 cells per strain per replicate. The indicated samples were compared using a one-sided t-test. (B) The distribution of Rad52 focus intensity is plotted for the indicated strains, after treatment with 50 *μ*g/ml phleomycin for 30 minutes (Phleo). The width of the box indicates the number of foci with a given intensity, and the bar indicates the mean. N=30 for all samples. The distributions were compared using the Wilcoxon rank sum test. (C) The direct repeat recombination rate was measured for the indicated strains. Each assay was a fluctuation test of nine cultures. The bars show the means of 10 replicates. (D) Schematic of the strain used to measure BIR. A DSB is induced by expression of the HO endonuclease. The right arm of chromosome III invades the left arm, and replication restores a functional *URA3* gene. (E) BIR was quantified for wild type and *mte1*Δ strains. N ranged from 3 to 6. The bars show the means of the replicates, and strains were compared using a t-test.

### Mte1 suppresses break-induced replication

Mph1 suppresses break-induced replication during doublestrand break repair (Luke-Glaser and Luke 2012; Stafa *et al*. 2014). Given the physical and genetic interactions between Mte1 and Mph1 that our work has revealed, we tested whether Mte1 also plays a role in suppressing BIR. We induced a DSB in strains carrying a modified chromosome V with a truncated *ura3* allele adjacent to an HO endonuclease site. Upon induction of the double-strand break, the truncated allele is repaired using donor sequences located on the other arm of chromosome V to yield Ura+ colonies (Figure 6D). In homologous BIR, where the sequences that recombine share 108 bp of homology, deletion of *mte1* results in increased BIR (Figure 6E), similar to deletion of *mph1* (Stafa *et al*. 2014). Thus, Mte1, like Mph1 is an important suppressor of break-induced replication and therefore a suppressor of loss of heterozygosity.

## Discussion

In response to DNA damage, most homologous recombination proteins are recruited to the sites of double-strand DNA breaks. Among them, Rad52 is a key recombination protein and the Rad52 focus is considered to be a sensitive indicator of DNA repair (Alvaro *et al*. 2007; Lisby *et al*. 2003 2001). We identified 29 proteins that localize to Rad52 foci in response to DNA damage. Among them, we identified a role for *YGR042W*/*MTE1* in DNA double-strand break repair. Similar to many DNA repair proteins, Mte1 forms nuclear foci in response to double-strand breaks and Mte1 foci only form when the DNA helicase Mph1 is present. Mte1 forms protein complexes with Mph1, and both proteins are recruited to the chromatin flanking double-strand DNA breaks *in vivo*. In the absence of Mte1 the Rad52 repair centers accumulate, and Mte1 is important for suppressing break-induced replication. Together our data indicate that the function of Mph1 in recombination repair of double-strand breaks requires Mte1.

### How does Mte1 impact Mph1 function?

Mte1 and Mph1 appear to be members of a constitutive complex. The interaction between the two proteins, whether direct or indirect, was readily detected by co-immunoprecipitation of either protein even in the absence of DNA damage. Mte1 is important for Mph1 nuclear focus formation, and more importantly, for the recruitment of Mph1 to double-strand breaks *in vivo*. These data suggest that Mph1 functions as part of a protein complex containing Mte1. Consistent with this notion, deletion of *MTE1* conferred sensitivity to phleomycin that was similar to that conferred by deletion of *MPH1*, and the *mte1*Δ *mph1*Δ double mutant was no more sensitive, indicating that the genes function in the same genetic pathway for phleomycin resistance.

Our data suggest that Mte1 is not simply a structural component of Mph1 complexes, as Mte1 appears to have little effect on Mph1 stability *in vivo*. Mte1 could presumably play a role in targeting Mph1 to specific substrates *in vivo*. Such a role would be consistent with our findings that Mph1 nuclear foci and the recruitment or retention of Mph1 at double-strand breaks is compromised when Mte1 is absent. Mph1 suppresses cross-overs and break-induced replication by unwinding D-loop recombination intermediates (Mazon and Symington 2013; Prakash *et al*. 2009; Stafa *et al*. 2014). We find that Mte1 suppresses break-induced replication much like Mph1, thus it is also possible that Mte1 facilitates some aspect of Mph1 catalysis. Mte1 lacks obvious catalytic domains, and purified Mph1 is capable of unwinding D-loops and extended D-loops *in vitro* in the absence of Mte1 (Prakash *et al*. 2009; Sebesta *et al*. 2011; Sun *et al*. 2008). Nonetheless, it will be of great interest to determine whether Mte1 modulates Mph1 activity *in vitro*, as it appears that *in vivo* Mph1 is normally assembled into complexes that contain Mte1.

### Orthologues of Mte1

Mte1 has readily identifiable orthologues in other yeasts, including *Kluveromyces, Candida, Pichia*, and *Ashbya* species. Mte1 appears to be an orthologue of the *Schizosaccharomyces pombe* Dbl2 protein (Yu *et al*. 2013). Dbl2 colocalizes with the fission yeast Rad52, and with double-strand breaks, and is important for nuclear focus formation by Fml1, the fission yeast orthologue of Mph1 (Yu *et al*. 2013). Dbl2 does not have a clear role in Fml1 inhibition of cross-overs or inhibition of BIR as of yet, so it is not known if Dbl2 plays a functional role similar to Mte1. Mte1 contains a domain of unknown function, DUF2439, that is found in the human ZGRF1 protein. The DUF2439 domain is also found in Dbl2 (Yu *et al*. 2013), but does not appear to be important for DNA damage resistance or for nuclear focus formation. Further, ZGRF1 is likely membrane-anchored and so might not be a true orthologue of Mte1. Nonetheless, as several lines of evidence suggest that Mph1 is an orthologue of the human FANCM protein (Whitby 2010), our evidence that Mph1 functions in concert with an important cofactor suggests that FANCM might also require a protein partner for effective function *in vivo*.

## Acknowledgments

We thank members of the laboratories of Brenda Andrews and Charlie Boone for assistance with microscopy, and Tobit Glenhaber and Linus Glenhaber for assistance in constructing *mte1* deletion strains to measure BIR efficiency. We also thank Lorraine Symington and Daniel Durocher for providing strains. This work was supported by grants from the Canadian Cancer Society Research Institute (Impact grant 702310 to GWB), the Cancer Research Society (to GWB), the Natural Sciences and Engineering Research Council of Canada (Discovery Grant grant 327612 to ZZ), and the National Institutes of Health (GM76020 to JEH).

## Author contributions

AY: Designed and carried out the experiments, wrote the paper, edited the paper. TK: Analyzed ChIP-seq data, edited the paper. RA: Performed and analyzed BIR assays. SM: Performed recombination assays, edited the paper. JO: Performed recombination analysis, constructed strains. JEH: Edited the paper. ZZ: Analyzed ChIP-seq data, edited the paper. GWB: Designed the experiments, wrote the paper, edited the paper.

---

**Figure S1.**
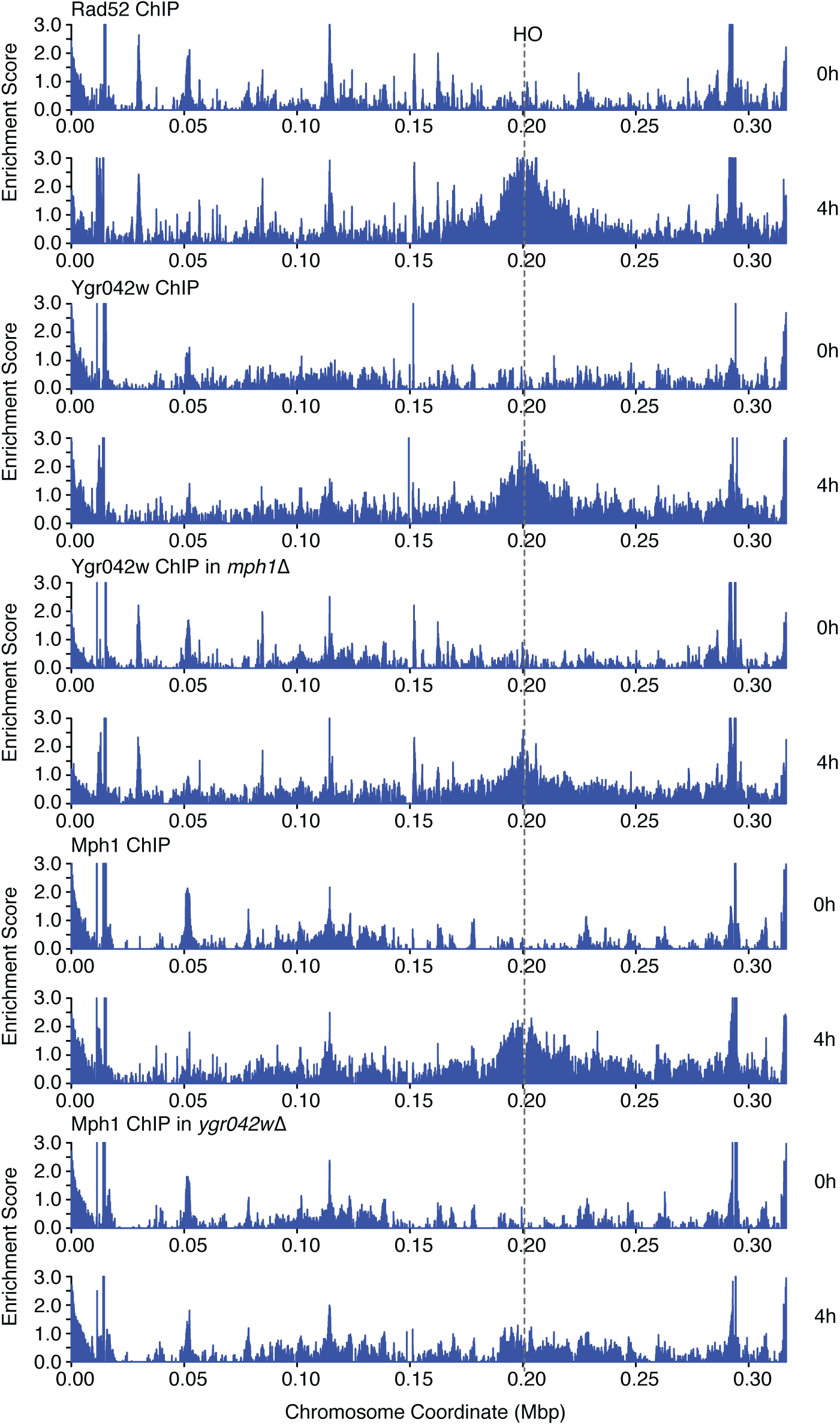
Mte1 and Mph1 are recruited to double-strand DNA breaks. ChIP-seq analysis was performed on *RAD52-FLAG, MTEI-FLAG, MTEI-FLAG Mph1*Δ, *MPHI-FLAG*, and *MPHI-FLAG mte1*Δ cells at 0h and 4h following the induction of a specific double-strand break at the MAT locus by the HO endonuclease. ChIP enrichment scores representing the log2 immunoprecipitate to input ratio are plotted across chromosome III for each time point. The position of the HO cut site is indicated.

**Table S1.**
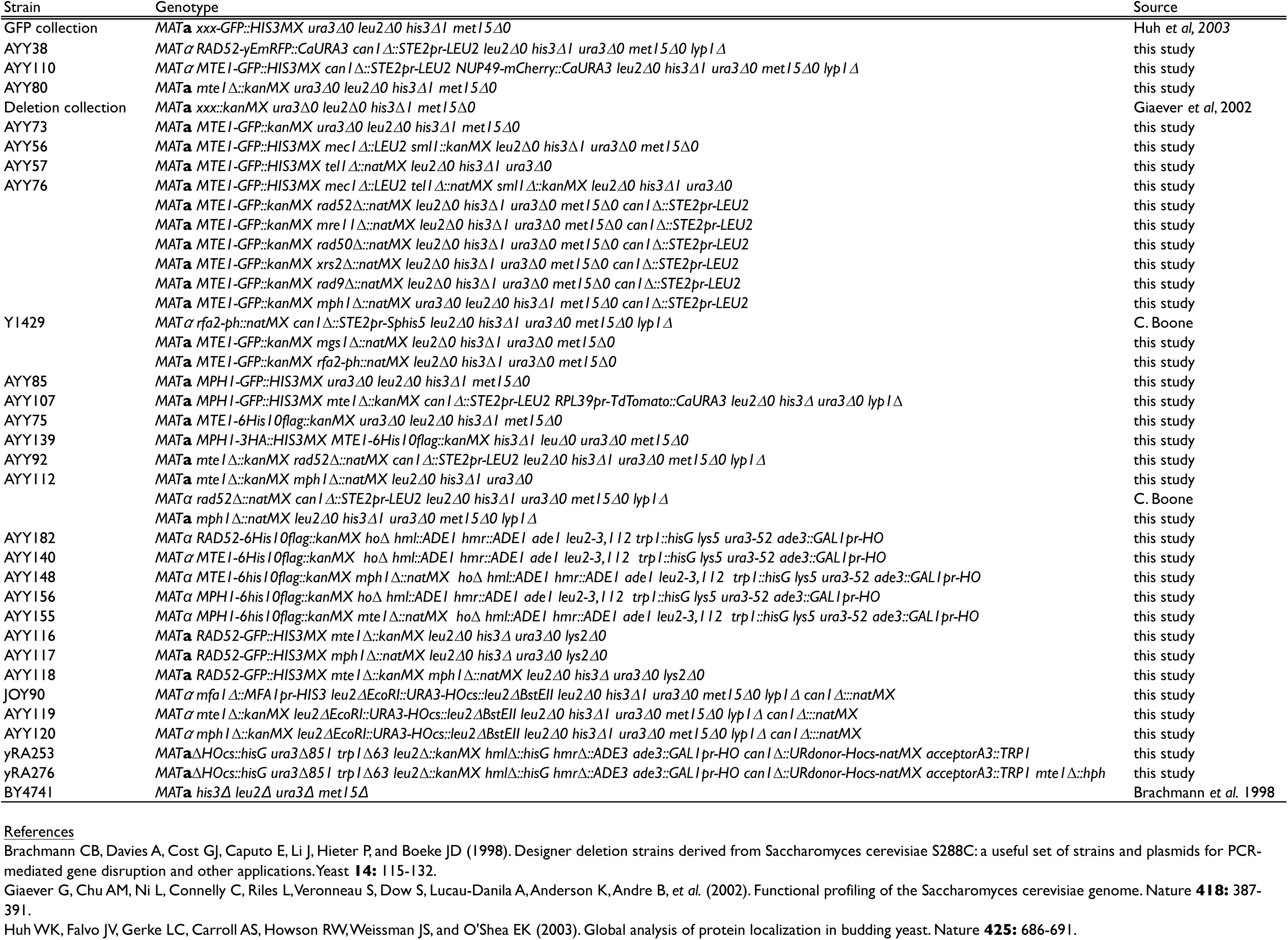
Yeast Strains used in this study.

**Table S2.**
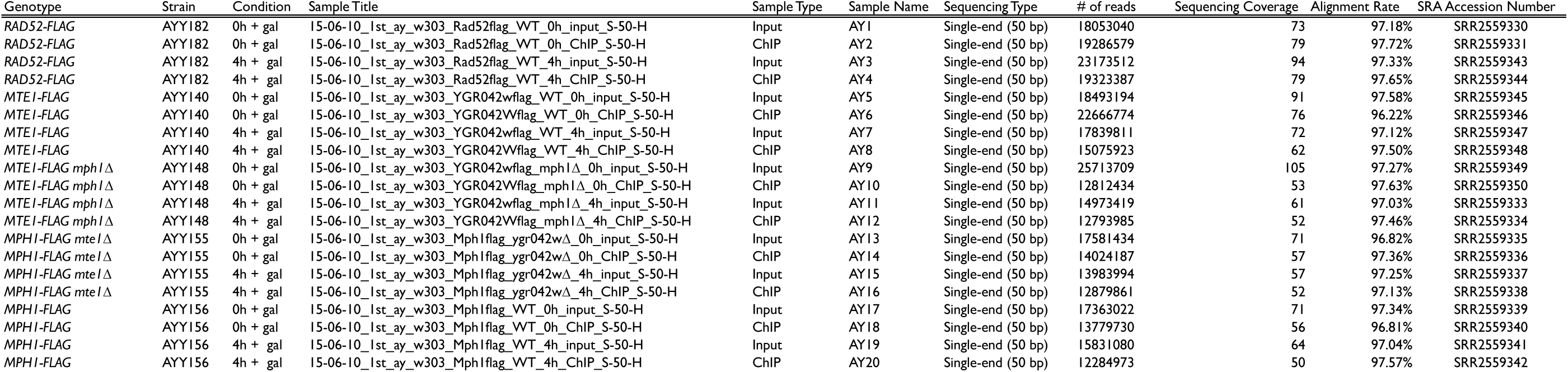
ChIp-seq statistics

**Table S3.** DFP fusion proteins that co-localize with Rad52 during phleomycin treatment (percent of foci)

| GFP-protein | ORF | % of GFP-Xxx foci that overlap Rad52 foci | % of Rad52 foci that overlap GFP-Xxx foci | # of Rad52-RFP foci | # of Xxx-GFP foci | Number of cells | Previously described co-localization with Rad52 |
| --- | --- | --- | --- | --- | --- | --- | --- |
| Srs2 | YJL092W | 78.9 | 29.4 | 51 | 19 | 201 | yes<sup>a</sup> |
| Rad59 | YDL059C | 54.5 | 30.0 | 20 | 11 | 147 | yes<sup>b</sup> |
| Ygr042w | YGR042W | 54.4 | 46.8 | 158 | 136 | 889 |  |
| Rad57 | YDR004W | 53.3 | 9.8 | 82 | 15 | 216 |  |
| Rad55 | YDR076W | 52.6 | 23.3 | 43 | 19 | 315 |  |
| Ddc2 | YDR499W | 47.1 | 71.4 | 91 | 138 | 197 | yes<sup>b</sup> |
| Mgs1 | YNL218W | 43.2 | 35.6 | 90 | 74 | 482 |  |
| Dna2 | YHR164C | 40.7 | 61.2 | 139 | 209 | 641 | yes<sup>b</sup> |
| Top3 | YLR234W | 37.5 | 12.5 | 24 | 8 | 185 |  |
| Rad53 | YPL153C | 34.8 | 25.0 | 96 | 69 | 329 |  |
| Rfa1 | YAR007C | 34.5 | 78.4 | 37 | 84 | 103 | yes<sup>c</sup> |
| Rad9 | YDR217C | 34.0 | 39.5 | 43 | 50 | 227 | yes<sup>b</sup> |
| Dpb11 | YJL090C | 31.3 | 57.7 | 26 | 48 | 102 | yes<sup>d</sup> |
| Rmi1 | YPL024W | 30.4 | 12.1 | 58 | 23 | 318 |  |
| Rad54 | YGL163C | 30.2 | 30.2 | 43 | 43 | 166 | yes<sup>b</sup> |
| Rfa2 | YNL312W | 27.4 | 65.4 | 26 | 62 | 59 | yes<sup>b</sup> |
| Sgs1 | YMR190C | 26.3 | 12.5 | 40 | 19 | 162 |  |
| Ddc1 | YPL194W | 24.3 | 71.4 | 35 | 103 | 118 | yes<sup>b</sup> |
| Rtt107 | YHR154W | 18.8 | 26.1 | 23 | 32 | 131 |  |
| Rad24 | YER173W | 17.6 | 2.0 | 153 | 17 | 430 |  |
| Rdh54 | YBR073W | 16.3 | 31.3 | 67 | 129 | 402 |  |
| Slx4 | YLR135W | 14.0 | 30.8 | 26 | 57 | 169 |  |
| Rtt101 | YJL047C | 12.5 | 4.5 | 111 | 40 | 563 |  |
| Mms21 | YEL091C | 9.3 | 8.0 | 50 | 43 | 258 |  |
| Mre11 | YMR224C | 9.1 | 7.7 | 39 | 33 | 123 | yes<sup>b</sup> |
| Slx8 | YER116C | 8.0 | 3.1 | 64 | 25 | 281 |  |
| Rad5 | YLR032W | 7.0 | 5.0 | 80 | 57 | 676 |  |
| Rad50 | YNL250W | 4.0 | 3.6 | 55 | 50 | 244 |  |
| Csm1 | YCR086W | 1.7 | 1.6 | 63 | 60 | 269 |  |

a. Burgess, R.C., Lisby M., Altmannova V., Krejci L., Sung P., and Rothstein R, 2009 Localization of recombination proteins and Srs2 reveals anti-recombinase function in vivo. *The Journal of Cell Biology* 185(6):969-981
b. Lisby, M., J.H. Barlow, R.C. Burgess, and R. Rothstein, 2004 Choreography of the DNA Damage Response; Spatiotemporal Relationships among Checkpoint and Repair Proteins. *Cell* 118 (6):699-713
c. Zhu, Z., Chung, W.H., Shim, E.Y., Lee, S.E., Ira, G., 2008 Sgs1 helicase and two nucleases Dna2 and Exo1 resect DNA double-strand break ends. *Cell* 134(6): 981-994
d. Germann, S.M., Oestergaard, V.H., Haas, C., Salis, P., Motegi, A., and Lisby, M, 2011 Dpb11/TopBP1 plays distinct roles in DNA replication, checkpoint response and homologous recombination. *DNA Repair (Amst)* 10(2):210-224
Pink shading indicates Xxx-GFP co-localizations less frequent than Mre11-GFP, and so should be regarded with caution.

**Table S4.** GFP fusion proteins that co-localize with Rad52 during phleomycin treatment (percent of cells)

| GFP-protein | ORF | Number of cells | Percent of cells with Rad52-RFP foci | Percent of cells with Xxx-GFP foci | Percent of cells with co-localized foci |
| --- | --- | --- | --- | --- | --- |
| Ddc1 | YPL194W | 197 | 8.6 | 24.9 | 27.4 |
| Rfa1 | YAR007C | 103 | 7.8 | 36.9 | 24.3 |
| Rfa2 | YNL312W | 59 | 13.6 | 39.0 | 22.0 |
| Ddc2 | YDR499W | 118 | 5.9 | 39.0 | 19.5 |
| Dpb11 | YJL090C | 102 | 5.9 | 28.4 | 13.7 |
| Dna2 | YHR164C | 641 | 5.0 | 14.2 | 12.5 |
| Ygr042w | YGR042W | 889 | 9.0 | 6.0 | 7.5 |
| Rad9 | YDR217C | 227 | 11.0 | 13.2 | 7.5 |
| Srs2 | YJL092W | 201 | 16.4 | 2.0 | 7.5 |
| Rad53 | YPL153C | 329 | 17.9 | 14.9 | 6.7 |
| Rad54 | YGL163C | 166 | 16.9 | 14.5 | 6.6 |
| Mgs1 | YNL218W | 482 | 10.6 | 7.3 | 5.8 |
| Rdh54 | YBR073W | 402 | 10.2 | 23.4 | 5.0 |
| Rtt107 | YHR154W | 131 | 13.0 | 16.0 | 4.6 |
| Slx4 | YLR135W | 169 | 10.7 | 23.1 | 4.1 |
| Rad59 | YDL059C | 147 | 7.5 | 3.4 | 4.1 |
| Rad57 | YDR004W | 216 | 27.8 | 2.8 | 3.7 |
| Rad55 | YDR076W | 315 | 9.8 | 2.9 | 3.2 |
| Top3 | YLR234W | 162 | 19.8 | 8.6 | 3.1 |
| Mre11 | YMR224C | 123 | 23.6 | 21.1 | 2.4 |
| Rmi1 | YPL024W | 318 | 14.2 | 4.1 | 2.2 |
| Sgs1 | YMR190C | 185 | 10.8 | 2.7 | 1.6 |
| Mms21 | YEL091C | 258 | 17.4 | 15.1 | 1.6 |
| Rtt101 | YJL047C | 563 | 18.7 | 6.0 | 0.9 |
| Rad50 | YNL250W | 244 | 19.3 | 16.4 | 0.8 |
| Slx8 | YER116C | 281 | 19.9 | 7.1 | 0.7 |
| Rad5 | YLR032W | 676 | 10.1 | 7.0 | 0.6 |
| Rad24 | YER173W | 430 | 28.4 | 3.3 | 0.5 |
| Csm1 | YCR086W | 269 | 21.9 | 23.8 | 0.4 |

**Table S5.**
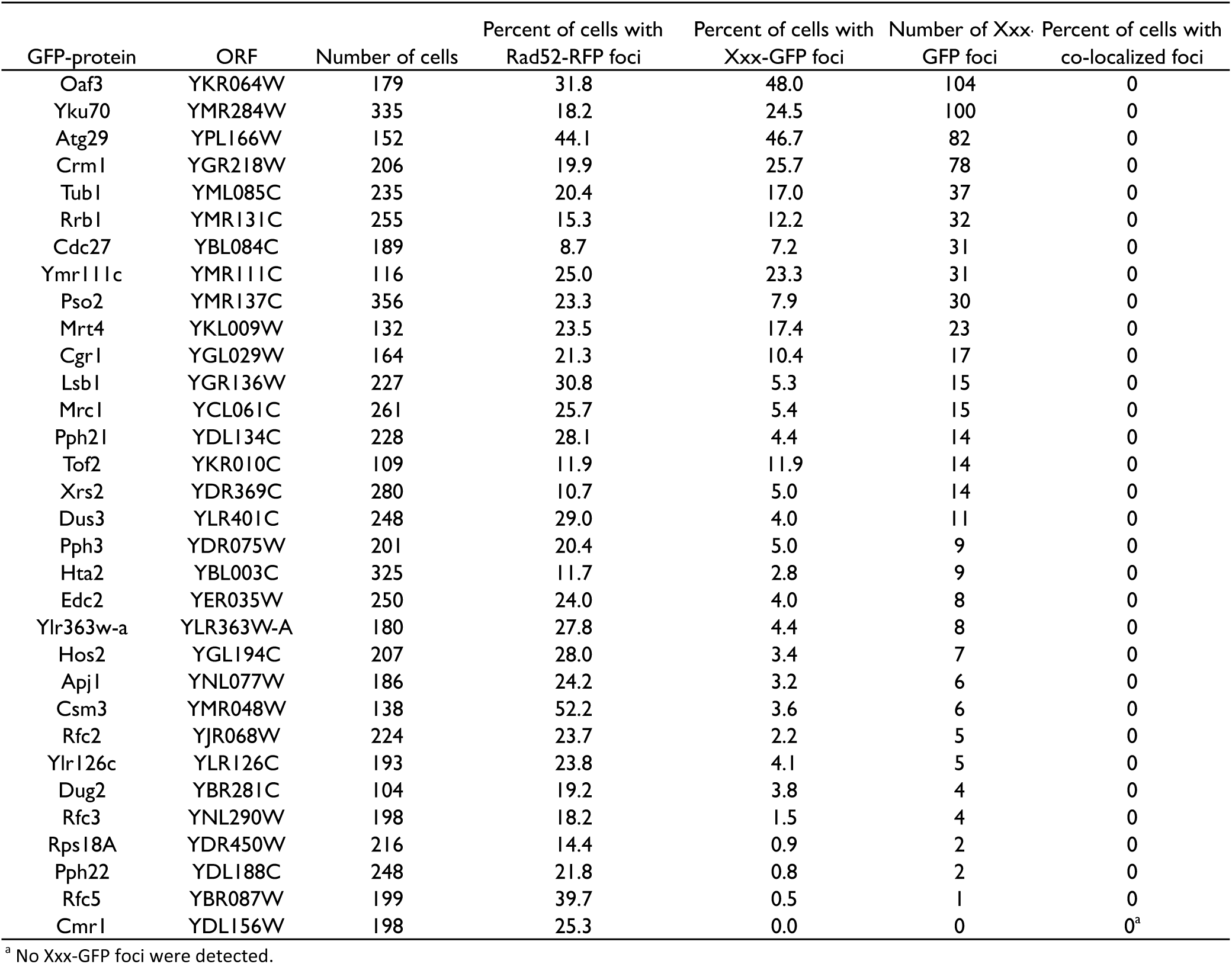
GFP fusion proteins that do not co-localize with Rad52 during phleomycin treatment (percent of cells)

**Table S6.**
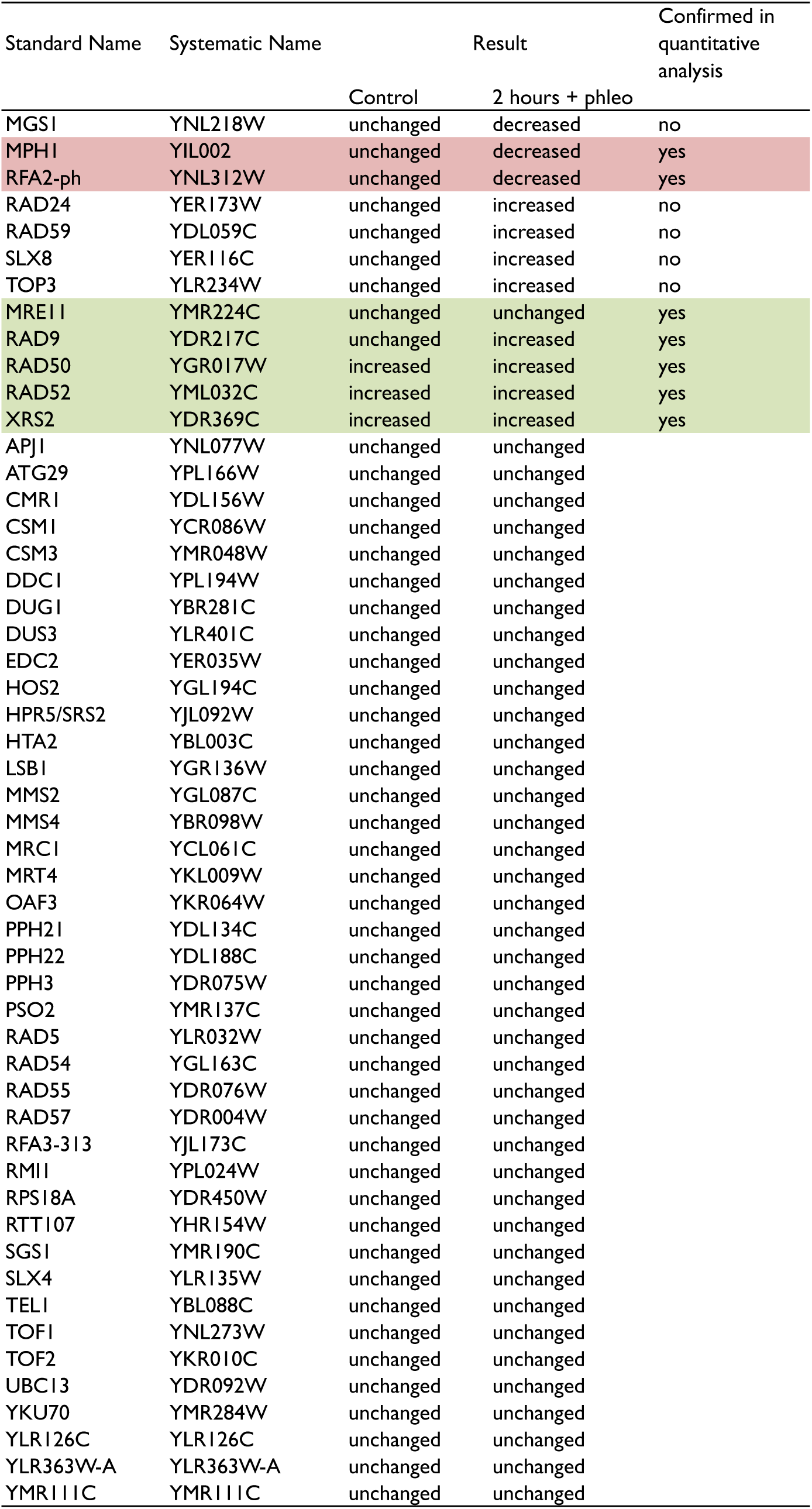
Mutants that affect Mte1-GFP focus formation

